# From Sequence to Significance: A Thorough Investigation of the Distinctive Genome Traits Uncovered in *C. werkmanii* strain NIB003

**DOI:** 10.1101/2023.11.21.568014

**Authors:** Mohammad Uzzal Hossain, Neyamat Khan Tanvir, Zeshan Mahmud Chowdhury, A.B.Z Naimur Rahman, Md. Shahadat Hossain, Shajib Dey, Arittra Bhattacharjee, Ishtiaqe Ahmed, Abu Hashem, Keshob Chandra Das, Chaman Ara Keya, Md. Salimullah

**Affiliations:** Bioinformatics Division, National Institute of Biotechnology, Ganakbari, Ashulia, Savar, Dhaka-1349, Bangladesh; Dept. of Biotechnology and Genetic Engineering, Jahangirnagar University, Savar, Bangladesh; Dept. of Microbiology, Noakhali Science and Technology, Noakhali, Bangladesh; Dept. of Biotechnology and Genetic Engineering, Noakhali Science and Technology, Noakhali, Bangladesh; Dept. of Mathematics and Natural Sciences, BRAC University, Dhaka, Bangladesh; Microbial Biotechnology Division, National Institute of Biotechnology, Ganakbari, Ashulia, Savar, Dhaka-1349, Bangladesh; Department of Biochemistry and Microbiology, North South University, Bashundhara, Dhaka-1229, Bangladesh

**Keywords:** genome, pathways, molecular, mutation, biological

## Abstract

*Citrobacter werkmanii* (*C. werkmanii*) is an opportunistic bacterium that has received very little investigation in Bangladesh, even though it is thought to be capable of causing diseases including diarrhea and urinary tract infections. Therefore, the present study focuses on the exploration of genomic features, including antibiotic resistance genes, virulent genes, and their distinctive nature in the pathogenesis of diseases. A complete genome was sequenced for the first time in Bangladesh after the morphological, biochemical and molecular identification of this bacteria. A total of 47,65,861 nucleotide base pairs makes up the genome, which contains 4669 genes, and many of the protein-coding genes can be classified into various ontological frameworks, including COG and KEGG pathways. Notably, *C. werkmanii* has the capability to have interactions with *Escherichia coli, Salmonella,* and *Klebsiella* while they possess the same domains and motifs, indicating a causal role in the development of diarrhea. Furthermore, the Bangladeshi isolate of *C. werkmaii* has observed genetic heterogeneity when compared to other strains of *C. werkmanii: C. werkmanii FDAARGOS_616, C. werkmanii LCU-V21*, and *C. werkmanii BF-6*. This identified genetic variation of sequenced *C. werkmanii* could drive the functional aberration in body homeostasis of bacteria. Later, molecular dynamics studies have proven the impact of their genetic variation on the protein structure, as indicated by the SASA, RMSD, Rg, and RMSF values. Finally, the findings of *C. werkmanii* from the sequenced genome might provide us with a better understanding of *C. werkmanii* and their possible role as opportunistic bacteria in diseases including diarrhea and urinary tract infections.

## Introduction

*Citrobacter* is becoming one of the most potent disease-causing agents worldwide due to its different transmission routes to humans, such as spoiled food, hospital tools and equipment, and fecal to oral transmission. *C. werkmanii* has been proven to be a causative agent of community-attained infections among neonates and immunodeficient patients (Kus, 2014; Ong et al., 2021).Neonatal meningitis and brain abscesses have been linked to *Citrobacter koseri*, whereas *Citrobacter freundii* causes neonatal meningitis and septicemia (Murray et al., 2010). Additionally, *Citrobacter* can cause diseases like those manifested by the urinary tract, the bloodstream, and the respiratory system, like lung abscess, bronchitis, and pneumonia (Ranjan & Ranjan, 2013). More significantly, wound infections, UTIs, and bacteremia have all been associated with *C. werkmanii* in the past (Madhumati et al., 2015; Mohanty et al., 2007). Although some strains can cause urinary tract infections, septic shock, and meningitis in infants, they are usually not the predominant cause of sickness. In general, Citrobacter species are not very dangerous, but studies in North America found that they were responsible for 4–5% of all nosocomial infections (Janda & Abbott, 2021). The gastrointestinal systems of humans and animals are just two of the many places where Citrobacter species can be found. These Citrobacter species can also be located in food, soil and as well as in water (Rogers et al., 2015). The *C. werkmanii AK-8* strain, separated from a stool sample in a hospital setting, exhibits the presence of antibiotic resistance genes, as confirmed by its complete genomic analysis (Parvez et al., 2020). Through the investigation of the AK-8 genome, some researchers discovered the presence of the two-component system of BaeSR, which controls the synthesis of efflux proteins (multidrug). The regulation of virulence was seen to be influenced by several factors, such as CpxRA, ZraRS etc. Additionally, it was discovered that Bar-UvrY has a role in controlling flagellum synthesis, the metabolism of carbon, and the formation of biofilm. The genome of *C. werkmanii AK-8* harbors a set of 20-21 chemoreceptors, which play a crucial role in facilitating colonization and pathogenicity (Parvez et al., 2020).

Their results highlight the importance of this bacterium as a widespread pathogen in the healthcare setting. *Citrobacter* bacterial species was recently known to be the second most frequent urinary pathogen in a survey done in Nepal. Additionally, this species was verified to be the third causative pathogen of urinary tract infections in India, accounting for up to 9.3%-9.4% of all isolates in a study involving urine samples from 4125 patients from January 2009 to December 2010 (Metri et al., 2013). *Citrobacter amalonaticus* was also found in 9.70% of patients in another study done in Northern India, where around 240 individuals tested positive for Citrobacter species (Sami et al., 2017).

Currently, there is a lack of studies undertaken in Bangladesh regarding the infection caused by *C. werkmanii*. Furthermore, there is an absence of comprehensive documentation on the whole genome of *C. werkmanii* within the country. The absence of genomic insights of this pathogenic bacterium is the reason for this. Multiple research findings indicate that C. werkmanii potentially exerts a substantial influence on the prevalence of diarrheal disease within the context of Bangladesh (Izquierdo et al., 2022).With the advancement of technology such as sequencing facilities (Sanger sequencing, Illumina sequencing), extensive bioinformatics tools, algorithms, and system biology approaches, we can investigate the genome and proteome of this bacteria and mitigate the problems caused by *C. werkmanii*.

In this research project, whole genome sequencing of *C. werkmanii NIB003* was done and searched for the similarities and differences with other strains of Citrobacter (*C. werkmanii FDAARGOS_616* and *C. werkmanii LCU-V21*) and also with other similar kinds of bacteria (*Escherichia, Salmonella,* and *klebsiella*). The increasing amount of research regarding multidrug-resistant (MDR) microorganisms indicates the urgent need to investigate innovative and more effective methods for addressing bacterial illnesses within the realm of public health. With the advancement of bioinformatics software and tools, this study will focus on the genomic insights of *C. werkmanii NIB003* and how this knowledge will enhance the public health sectors of Bangladesh.

## Materials and Methods

### Bacterial strain isolation and culture

Stool samples were collected and subsequently introduced into 9 ml of Tryptic soy broth. Our targeted sample was named after NIB003. The mixture was then subjected to incubation at a temperature of 37°C for a duration of 18 hours. Following the enrichment process, a single inoculum was subsequently spread over Xylose Lysine Deoxycholate agar (XLD agar) and incubated at a temperature of 37°C for a duration of 24 hours. Non-lactose fermenting microorganisms exhibiting translucent or colorless colonies were selected from Xylose Lysine Deoxycholate (XLD) agar plates. These colorless colonies were subsequently streaked onto fresh XLD agar plates and incubated at a temperature of 37°C for a duration of 24 hours. The suspected colonies of Citrobacter were selected from a selective medium and subsequently streaked onto nutrient agar plates. These plates were then incubated at a temperature of 37° C for a period of 18-24 hours to facilitate the biochemical characterization of the bacteria.

### Morphological and molecular characterization

Typical colonies were selected from Nutrient Agar plates and were subjected to slide preparation, Gram staining, microscopic observation, and biochemical tests. Pure colonies were identified by biochemical tests, triple sugar iron agar (TSI), Motility test, Urease test, Indole test, citrate test, and oxidase test. The morphology was examined using a microscope (Guzman et al., 2022). The extraction and purification of bacterial genomic DNA were performed using the Thermo Scientific GeneJET Genomic DNA Purification Kit and following their designated extraction technique. The bacterial universal primers 27F (5’-AGA GTT TCA TCT GGC TCA G-3’) and 1492R (5’-GGT TAC CTT GTT ACG ACT T-3’) were used to amplify the 16S rDNA gene in the genomic DNA of a high-yield, protease-producing bacterium. The DNA from this bacterium was used as a template for the amplification process. The polymerase chain reaction (PCR) was then performed by following the standard protocols. Afterwards, the PCR products were loaded onto the 3130 Genetic Analyzer (Applied Biosystems). The sequences were constructed automatically using Sequencing Analysis 5.3.1 software, developed by Applied Biosystems (Faja et al., 2019). The spliced sequences were subjected to comparison with the data available in the NCBI database. The CLC drug discovery software was utilized to study the standard bacteria that exhibited comparable genetic links (Stackebrandt & Goebel, 1994).

### Whole genome sequencing (WGS)

The bacterial isolates that were identified and validated were subjected to sequencing using the MiSeq system of the Illumina sequencing platform at the National Institute of Biotechnology’s center for next-generation sequencing facilities. Initially, the genomic DNA was fragmented into fragments ranging from 400 to 550 base pairs (bp) through the utilization of the M220 focused-ultrasonicator (Covaris Ltd. Brighton, UK). The sequencing library was prepared with the TruSeq DNA Library LT kit (Illumina, San Diego, CA, USA), following the established instructions provided by the manufacturer.

### WGS assembly and annotation

The quality control of raw sequencing data was conducted with the utilization of the Linux terminal and the FastQC command tools, followed by adapter sequence trimming using the Trimmomatic tool (Bolger et al., 2014). The NextTera adapter was utilized for the purpose of adapter sequence trimming. The generated parameters resulted in the production of two paired read files in the FASTQ format. After obtaining the paired-end reads, we utilized the Unicycler technique for conducting the genome assembly (Wick et al., 2017). Subsequently, the genome annotation command-line tools, predominantly Prokka (Seemann, 2014) were executed to annotate the assembled genome.

### Genomic exploration using computational analysis

To visualize and characterize the entire genome of NIB003, we used Artemis software (Carver et al., 2012). The contigs of the assembled genome were subjected to screening for virulence properties in the VFDB (Virulence Factor Database) (Chen et al., 2016). The current study utilized the entire antibiotic resistance database to find plausible gene candidates that are linked to antimicrobial resistance characteristics (Jia et al., 2017; B. Liu et al., 2019). This screening was performed using the ABRicate tool, version 1.0.1, with parameter cutoffs of 90% coverage and 95% nucleotide identity (Seemann T, n.d.). The ABRicate findings were subsequently visualized on a clustered heatmap, which illustrated the presence or lack of the virulence and antibiotic resistance gene profiles (Letunic & Bork, 2019).

For the visualization and exploration of common interacting genes in the complete genome of *C. werkmanii NIB003*, we used STRING (Szklarczyk et al., 2019) and Cytoscape (Shannon et al., 2003). Furthermore, we used CLC Drug Discovery software for multiple sequence alignment and the construction of a phylogenetic tree to illustrate the evolutionary relationship among *NIB003*, *Escherichia coli, Salmonella, and Klebsiella*. For motif and domain comparison, we used several bioinformatics tools, such as ScanProsite (de Castro et al., 2006), InterProScan (Quevillon et al., 2005), and Pfam (Finn et al., 2014). The process of gene function annotation involved the utilization of multiple online databases such as SwissProt (Bairoch & Apweiler, 2000), GO databases (Gene Ontology Consortium, 2006), COG (Tatusov et al., 1997), and KEGG (Kanehisa & Goto, 2000).

### Molecular Dynamic Simulation

The protein stability was scrutinized using the GROningen MAchine for Chemical Simulations (GROMACS, version 2022.3) (Kus, 2014) through a molecular dynamics simulation at 100 nanoseconds (ns). The simulation utilized the CHARMM36m force field. The TIP3 water model was employed in the construction of a water-box, with the boundaries of the box situated at a distance of 1 nm from the protein surface. The neutralization of the systems was achieved through the utilization of the requisite ions. Following the completion of energy minimization, isothermal-isochoric (NVT) equilibration, and isobaric (NPT) equilibration procedures, a molecular dynamic simulation with a duration of 100 nanoseconds was carried out. The simulation incorporated periodic boundary conditions and utilized a temporal integration step of 2 femtoseconds. The interval for capturing snapshots of the trajectory data was configured to be 100 picoseconds. Following the completion of the simulation, we employed the RMSD, RMSF, Rg, and SASA modules integrated within the GROMACS software to conduct an evaluation of the root mean square deviation (RMSD), root mean square fluctuation (RMSF), radius of gyration (Rg), and solvent accessible surface area (SASA). The ggplot2 package in RStudio was utilized to produce the visual representations for each of these research inquiries. The molecular dynamics (MD) simulations were performed using high-performance simulation workstations running on the Ubuntu 22.04.3 LTS operating system.

## Results

### Morphological and biochemical characterization of the bacteria

After culturing the targeted NIB003 bacteria onto Xylose Lysine Deoxycholate (XLD) agar medium, the bacteria were observed for further investigation. In this study, colonies that exhibited a black coloration or possessed red centers with black pigmentation were seen as indicative of positive results and a possible presence of *C. werkmanii* (**Fig. 1A**). Afterwards, selected bacteria were visualized under a microscope, which confirmed the presence *of C. werkmanii* (**Fig. 1B**). We further analyzed the several biochemical tests, including the absence of indole and urea production as well as the existence of bacterial motility observed in MIU agar medium. H2S production and gas were detected on triple sugar iron (TSI) medium. Daily recordings of all reactions were made, and the final readings were taken at the 72-hour mark. There were no observable alterations in responses after the 72-hour mark. The NIB003 sample was subjected to additional characterization based on its fermentation of sucrose, raffinose, ot-methylD-glucoside, and melibiose, as well as its capacity to use Simmons citrate. *C. werkmanii* conducted fermentation on the four aforementioned sugars and were citrate-negative (**Table 1 and Supplementary Fig. 1**).

**Figure 1.**
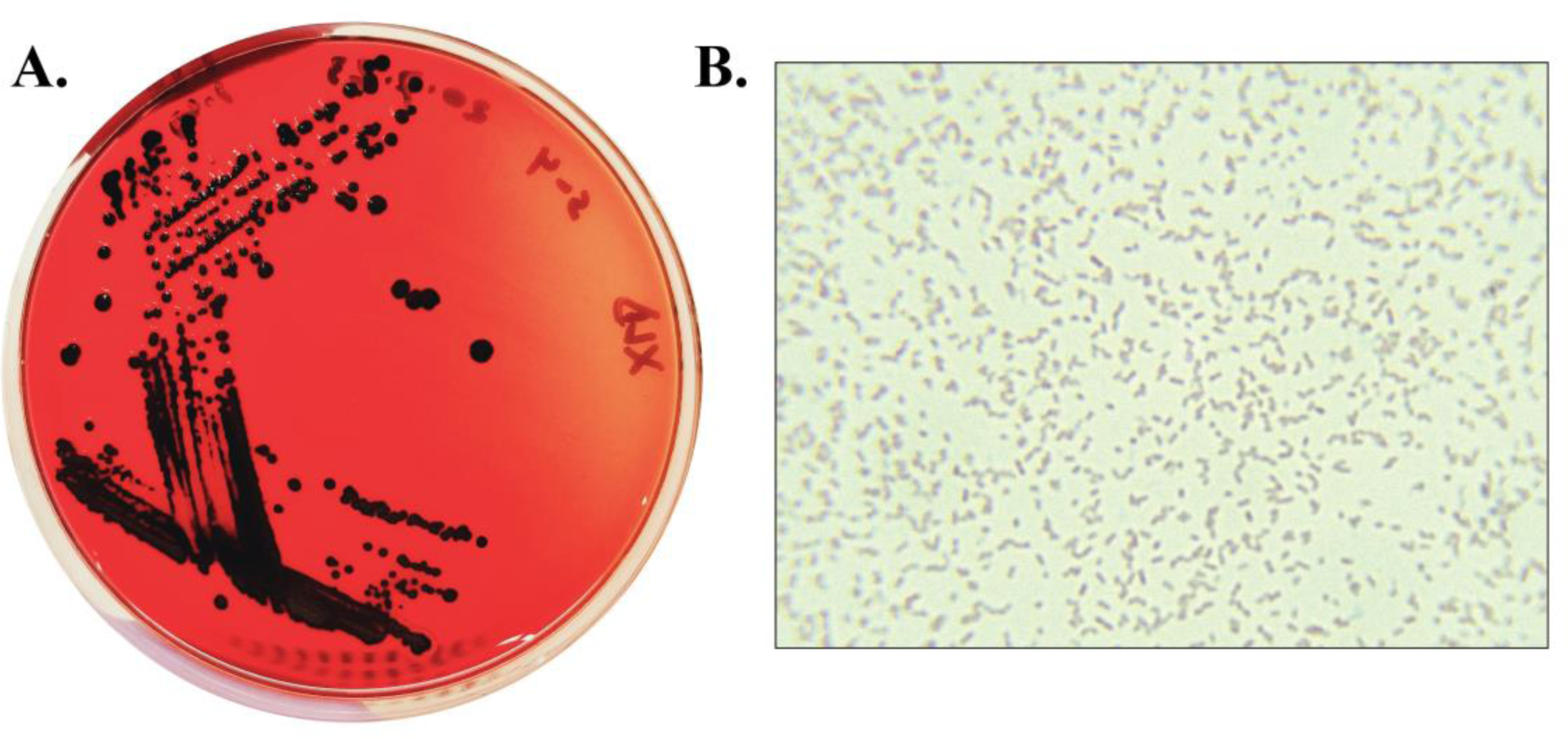
**A.** Culture of *Citrobacter werkmanii* on Xylose Lysine Deoxycholate (XLD) agar. **B.** Microscopic visualization of *Citrobacter werkmanii*

**Table 1.** Lists of biochemical tests for NIB003 sample.

| Sample ID | MIU medium |  |  | TSI medium |  |  |  | Citrate |
| --- | --- | --- | --- | --- | --- | --- | --- | --- |
|  | Motility | Indole | Urea | Slant | Butt | H<sub>2</sub>S | Gas |  |
| Citrobacter werkmanii | + | - | - | Alkaline<br>(Red) | Alkaline<br>(Red) | + | + | + |

### Molecular characterization of the bacteria

After agarose gel electrophoresis of PCR products, we determined the size of the PCR products ranged from 1.4kb-1.5 kb (**Fig. 2A**). We compared the 16S rRNA sequences of *C. werkmanii NIB003* and other closely related strains of the same genus. Then, we used the MEGA X software to make a phylogenetic tree with 100 bootstrap replicates using the Maximum Likelihood Tree and the Kimura 2-parameter methods (**Fig. 2B**).

**Figure 2.**
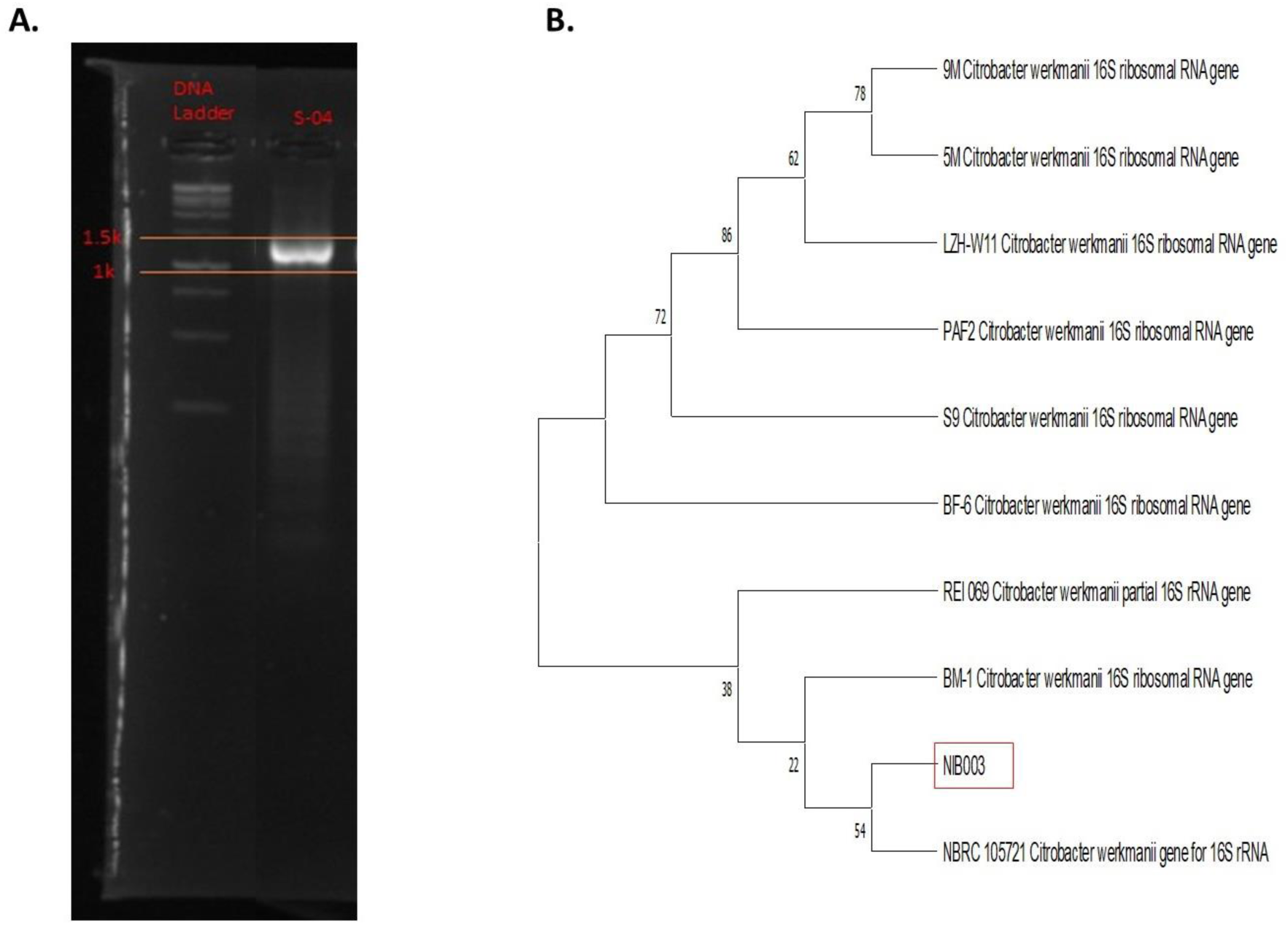
**A.** S-04: Band of *Citrobacter werkmanii*; PCR products visualization using agarose gel electrophoresis along with ethidium bromide, **B.** Phylogenetic tree based on 16S rRNA of *Citrobacter werkmanii*.

### Characterization of the complete genome

The *C. werkmanii NIB003* strain was subjected to whole genome sequencing, resulting in 40 contigs. The genome of this strain was determined to consist of a total of 47,65,861 nucleotide bases (**Fig. 3**). In addition, our study revealed the presence of 4437 coding sequences responsible for encoding numerous vital proteins involved in bacterial survival, motility, adherence to the host surface, and various other biological processes. These organisms possess a total of 4587 genes and a similar number of proteins. Among the proteins analyzed, there were 128 instances of miscellaneous RNA (misc_RNA). This bacterium contains a total of 4 ribosomal RNA (rRNA) molecules, 68 transfer RNA (tRNA) molecules, and 1 transfer-messenger RNA (tmRNA) molecule (**Table 2**). We’ve compared the Citrobacter species based on genomic differences (**Table 3)**. In the conducted study, it was observed that the genome size of *C. werkmanii NIB003* was determined to be 47,65861 base pairs (bp), a value comparable to that of *C. werkmanii FDAARGOS_616* (4,947,997 bp) *and C. werkmanii LCU-V21* (4,929,495 bp).

**Figure 3.**
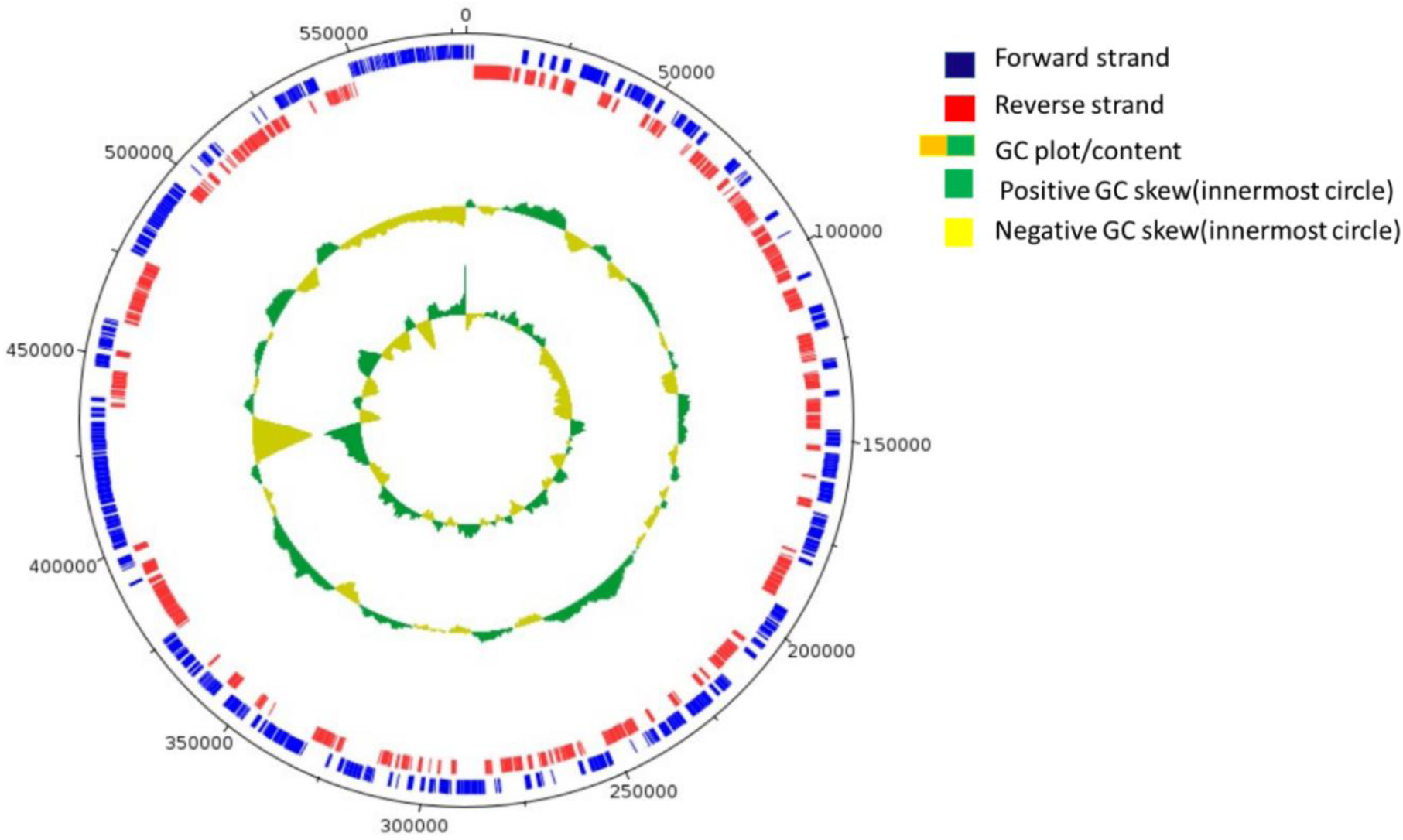
Genomic visualization of Citrobacter werkmanii NIB003 using ARTEMIS software. Visual depiction of genome, showing protein-coding regions and GC content. The circular representation consists of multiple concentric circles, each depicting different genomic features. Starting from the interior circle, circle 1 represents the GC skew, with positive GC skew indicated by the color green and negative GC skew represented by the color yellow. Moving outward, circle 2 displays the GC content/plot. Circle 3 shows genes on reverse strand(red); circle 4, genes on forward strand (blue); circle 5 visualizes genome size.

**Table 2.** General Features of *Citrobacter werkmanii NIB003* genome.

| Major features | Value |
| --- | --- |
| Genome size | 4765861 |
| Genes | 4587 |
| Proteins | 4465 |
| G+C contents | 52% |
| tRNA | 68 |
| rRNA | 4 |
| tmRNA | 1 |
| misc_RNA | 128 |

**Table 3.**
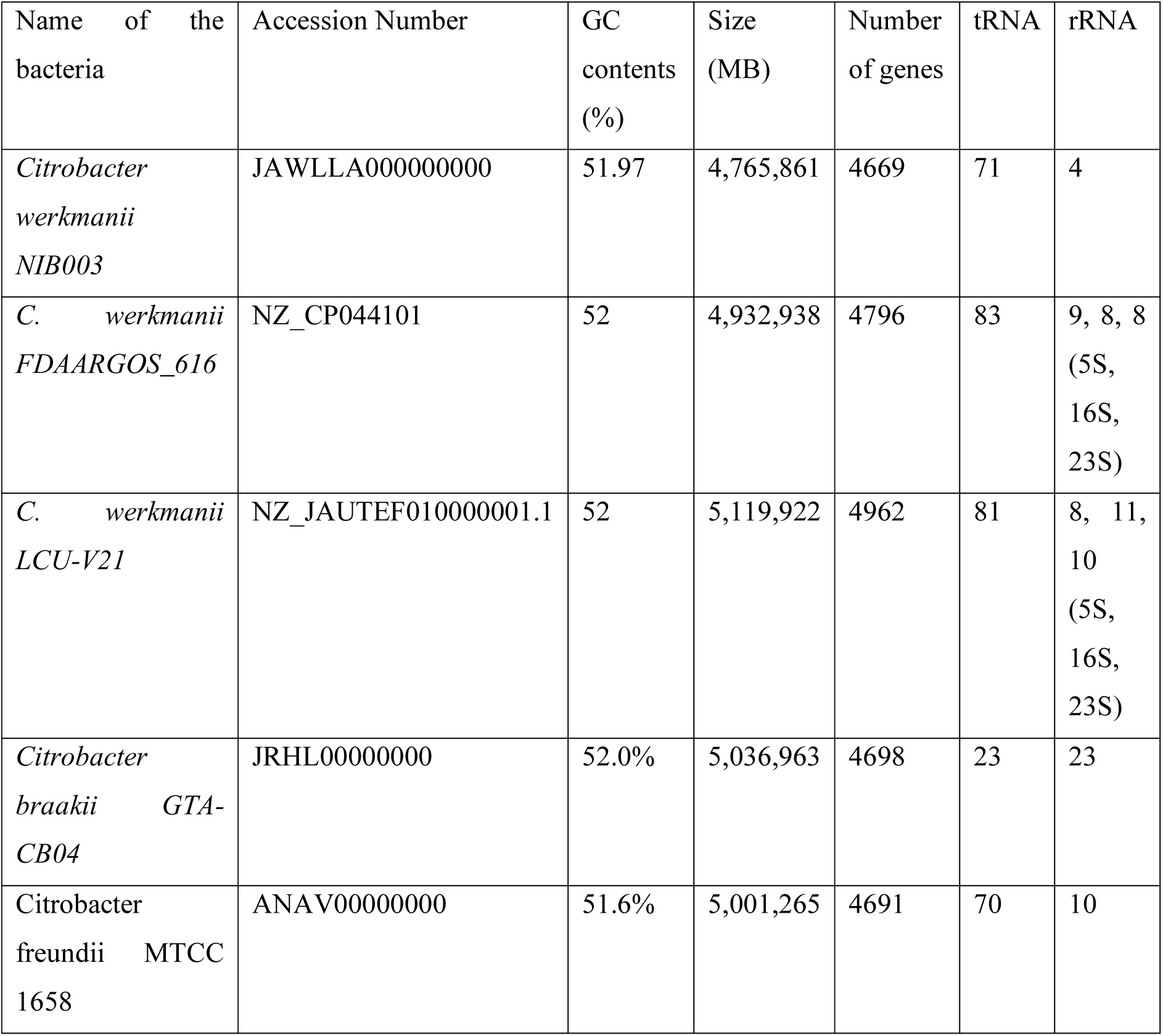
Comparison of *Citrobacter* species based on genomic differences.

### Characterization of the proteome of genome

#### Characterization of antibiotic resistance genes

The whole genome sequencing and subsequent bioinformatics analysis of *Citrobacter werkamnii NIB003* reveal two antibiotic resistance genes, such as blaCMY-98 and QnrB69_1, which are also found in *Citrobacter freundi*. The protein in the blaCMY-98 gene is Beta-lactamases. Beta-lactamase, also known as β-lactamases, are enzymatic proteins synthesized by bacteria. These enzymes provide resistance to a wide spectrum of beta-lactam drugs, including penicillins, cephalosporins, cephamycins, and carbapenems (specifically, ertapenem). The protein of QnrB69 is the Quinolone resistance pentapeptide repeat protein QnrB69. The gene has a sequence of repeated units known as pentapeptide repeats. The presence of these repetitive sequences was initially observed in numerous proteins of cyanobacteria, but subsequent investigations have revealed their occurrence in proteins of both bacterial and plant origins (Kieselbach et al., 1998). The precise activities of these repetitive elements remain elusive; nonetheless, empirical evidence has demonstrated that individuals within this gene family possess a common capacity to engage in interactions with DNA-binding proteins, including DNA gyrase. Moreover, the QnrB69 gene, which is present in *C. werkmanii NIB003*, plays a significant role in conferring resistance to the fluoroquinolones (Mérens et al., 2009).

#### Characterization of virulent genes

After whole genome sequencing of *C. werkmanii*, we identified a number of virulent genes. Virulence factors refer to the molecular components that facilitate the colonization of the host by bacteria at the cellular level. These factors can be classified as either secretory, membrane-associated, or cytosolic in nature. At locus PLHDHIGA_00948 of *C. werkmanii NIB003* genome, we found the toxin-antitoxin biofilm protein (TabA), which stimulates biofilm formation, hinders fimbria genes in biofilms, and may act in response to toxin-antitoxin systems. At locus PLHDHIGA_04053, we identified a protein called fliQ (flagellar biosynthetic protein), a protein involved in the biosynthesis of bacterial flagella that can facilitate motility towards host targets, biofilm formation to enhance bacterial survival, secretion of virulence factors, trigger both adaptive and innate immune defense mechanisms, and promote adherence and invasion. At PLHDHIGA_04108, a virulent protein called common pilus major fimbrillin subunit EcpA was found to play a dual role in early-stage biofilm development and host cell recognition.

### KEGG pathway of the proteome

In order to get insight into the internal metabolic pathways and the roles of gene products, the genes were correlated with their respective terms in the KEGG pathway database, such as ABC transporters, two component systems, purine metabolism, pyruvate metabolism, ribosome, flagellar assembly, quorum sensing, biofilm formation, pyrimidine metabolism, and glycolysis. In this respective pathway, ABC transporters route exhibited the greatest number of genes (170), signifying its prominence within this context. Conversely, the glycolysis pathway had the fewest genes (42), indicating its very limited involvement. Additionally, the flagellar assembly pathway, which contributes to virulence, encompassed 45 genes **(Fig. 4**).

**Figure 4.**
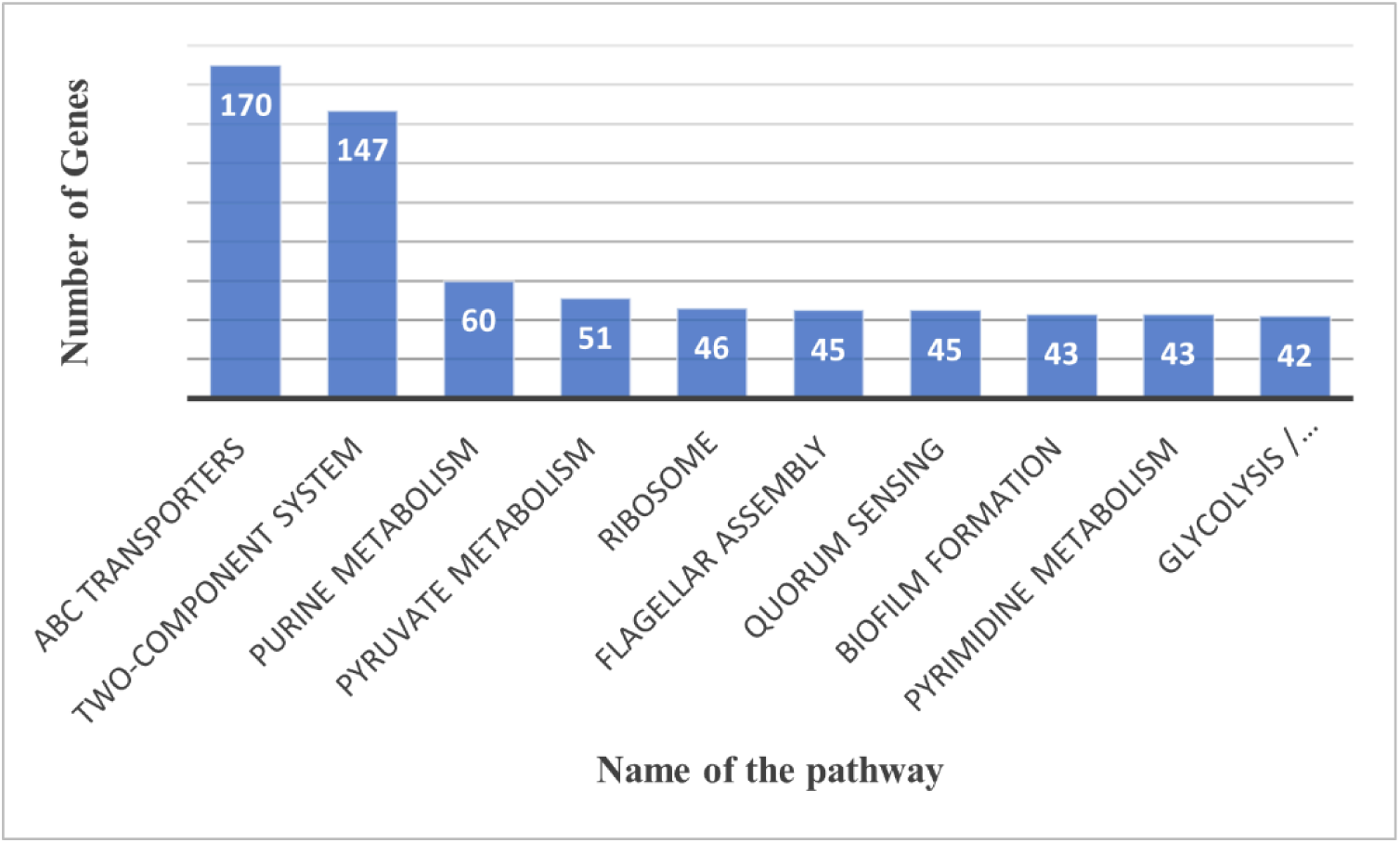
Top 10 pathway by *Citrobacter werkmanii* NIB003

### Finding the highly connected proteins from the proteome of *C. werkmanii NIB003*

A network of proteins is generated and shown using the software tools STRING and Cytoscape based on the provided protein list (**Fig. 5**). The 11 proteins of C. werkmanii NIB003 (PLHDHIGA_02079, PLHDHIGA_01872, PLHDHIGA_01946, PLHDHIGA_04004, PLHDHIGA_01430, PLHDHIGA_03404, PLHDHIGA_01118, PLHDHIGA_02178, PLHDHIGA_00433, PLHDHIGA_01431, PLHDHIGA_00328) were interacted with the proteins of three closest bacteria such as salmonella (STY2617,STY2109,asns,TY 1673,cysS,STY2280,StY2767,metG,serS,thrS,SBOV25871),Escherichia(Ecs1013,yfgM,Ecs3 376,Ecs2426,Ecs0978,Ecs3215,tyrs,Ecs0588,Ecs2820),Klebsiella(thrS_2,tyrs_2,AML34477. 1,hisG,prmB,sarS,KNC10201.1.AML38704.1,AEW60537.1,ygjH). In order to elucidate the evolutionary correlation between *C. werkmanii* NIB003 and other bacterial strains (*Salmonella, Escherichia coli, and Klebsiella*), a phylogenetic analysis was conducted using neighbor joining tree. The phylogenetic study demonstrated a significant evolutionary closeness between *C. werkmanii NIB003* and the bacterial genera *Escherichia, Klebsiella,* and *Salmonella* **(Supplementary Fig. 2).**

**Figure 5.**
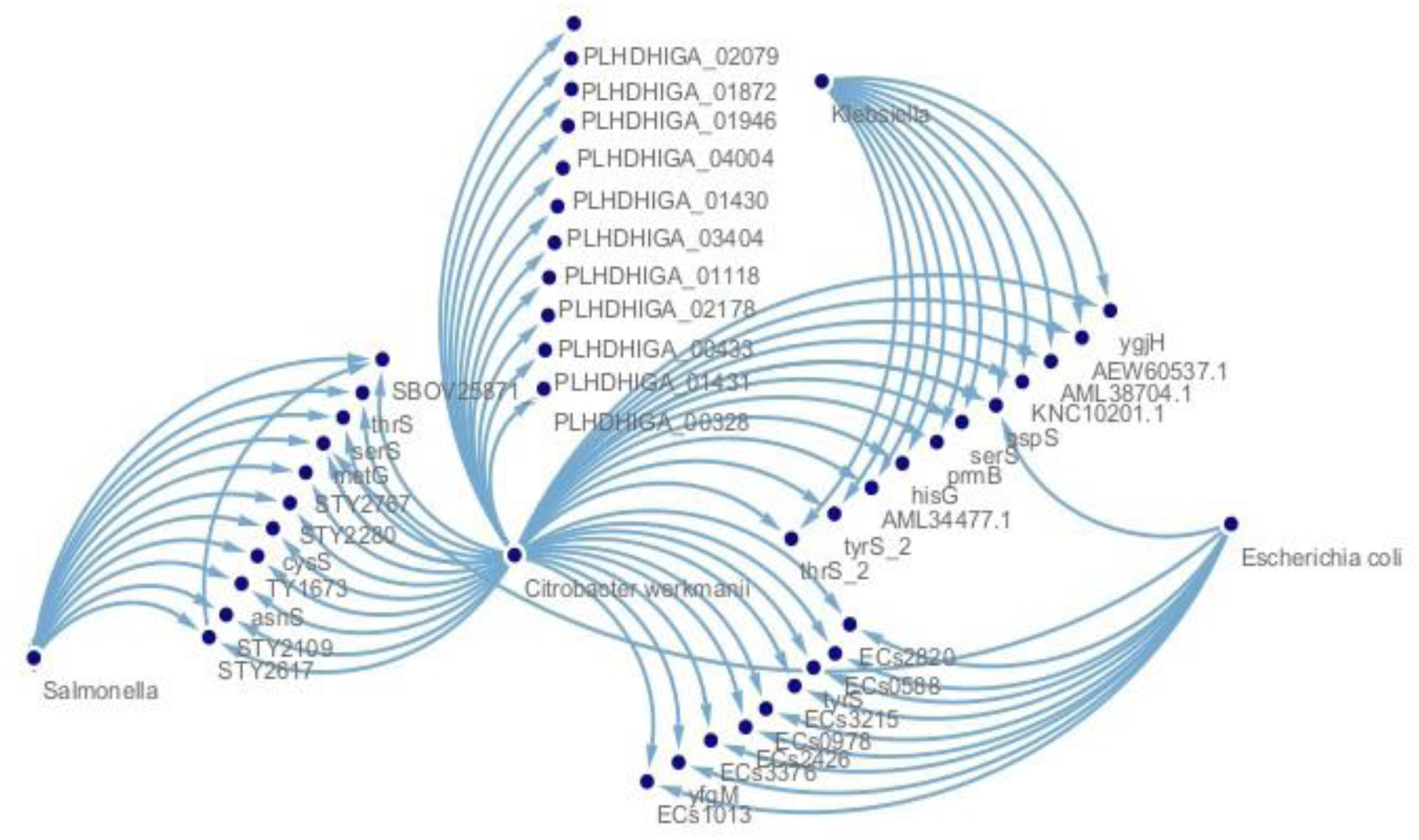
Interactions of different proteins of *C. werkmanii NIB003* with 3 closest bacteria (*Salmonella. Escherichia coli, Klebsiella*)

The gene products in the COG database were categorized into several clusters of orthologous groups. The three most prominent categories observed in the study were carbohydrate transport and metabolism, amino acid transport and metabolism, and transcription (**Table 4**).

**Table 4.** KEGG pathway of top 11 interacting proteins of *Citrobacter werkmanii NIB003*.

| Gene | KO Id | COG id | Pathway |
| --- | --- | --- | --- |
| PLHDHIGA_01430 | K01892 | COG0124 | Aminoacyl-tRNA biosynthesis |
| PLHDHIGA_01118 | K01874 | COG0143 | Selenocompound metabolism;<br>Aminoacyl-tRNA biosynthesis;<br>Metabolic pathways; |
| PLHDHIGA_01872 | K01893 | COG0017 | Aminoacyl-tRNA biosynthesis |
| PLHDHIGA_03404 | K01876 | COG0173 | Aminoacyl-tRNA biosynthesis |
| PLHDHIGA_00328 | K01875 | COG0172 | Aminoacyl-tRNA biosynthesis |
| PLHDHIGA_02178 | K01883 | COG0215 | Aminoacyl-tRNA biosynthesis |
| PLHDHIGA_02079 | K01866 | COG0162 | Aminoacyl-tRNA biosynthesis |
| PLHDHIGA_00433 | K00765 | COG0040 | Histidine metabolism;<br>Metabolic pathways; |
|  |  |  | Biosynthesis of secondary metabolites;<br>Biosynthesis of amino acids |
| PLHDHIGA_01946 | K01868 | COG0441 | Aminoacyl-tRNA biosynthesis |
| PLHDHIGA_04004 | K02493 | COG2890 | -- |

### Functional genomic differences of *C. werkmanii NIB003* with the closest bacteria

There are distinctive and similar motifs found in the above-mentioned four classes of bacteria (*C. werkmanii NIB003, Escherichia coli, Salmonella, and Klebsiella*). The presence of the tRNA binding domain motif (TRBD) may be observed in *C. werkmanii NIB003*, *E. coli CFT073*, and *Salmonella enteria diarizonae* **(Fig. 6 and Table 5).** The presence of ATP Phosphoribosyltransferase was also observed in the aforementioned bacterial species. Dam methylases were also identified in the aforementioned bacteria. The discovery of the S4 RNA-binding motif, which has the ability to recognize intricate three-dimensional characteristics within extensively folded RNA molecules, has been documented. The presence of the aminoacyl-transfer RNA synthetases class-II family, a widely observed pattern responsible for activating amino acids and facilitating their binding to certain tRNA molecules during the initial stage of protein production, has been identified in these bacterial organisms (**Table 5**). Moreover, we aligned *C. werkmanii NIB003* with other three bacteria (*Escherichia coli, Salmonella, and Klebsiella*) and found a nucleotide change (alanine instead of serine) in the aminoacyl-transfer RNA synthetases class-II motifs at locus PLHDHIGA_01872 of *C. werkmanii* genome, another change (glutamine instead of lysine) found in the same motifs as previously mentioned, at locus PLHDHIGA_03404 (**Supplementary Fig. 3**).

**Figure 6.**
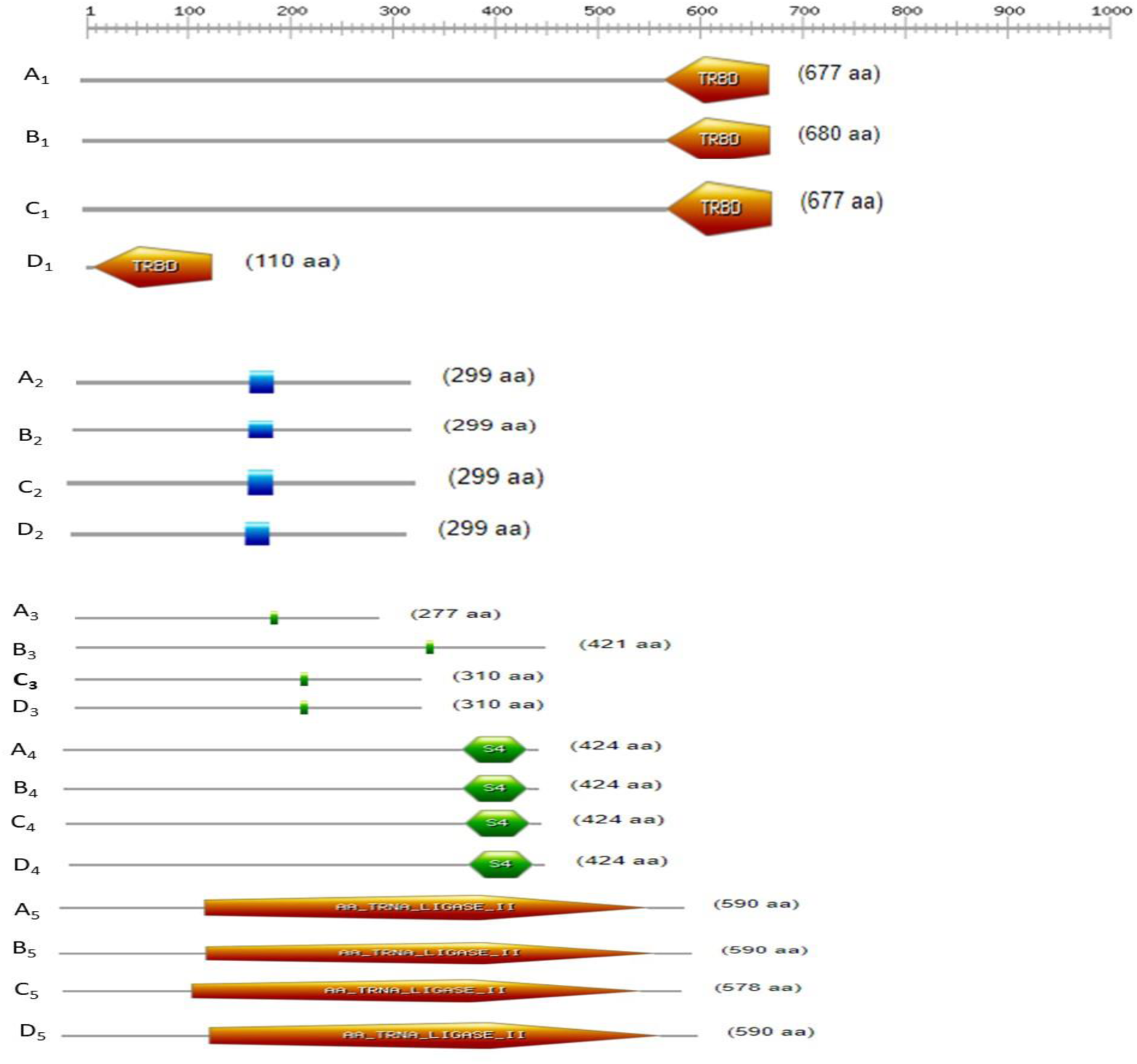
Comparison of motifs of *Citrobacter werkmanii NIB003* with three closest bacteria. **A_1_:** PLHDHIGA_01118; *Citrobacter werkmanii NIB003*; motif range (575 – 677), **B_1_:** METG; *Escherichia coli CFT073*; motif range (578-680), **C_1_:** METG; *Salmonella enterica diarizonae*; motif range (578-680), **D_1_:** YGJH; *Klebsiella quasipneumoniae*; motif range (8 – 110). **A_2_:** PLHDHIGA_00433; *Citrobacter werkmanii NIB003*; motif range (156-177) **B_2_**: ECS2820; *Escherichia coli O157H7 Sakai*; motif range (156 – 177), **C_2_:** STY2280; *Salmonella enterica Typhi*; motif range (156 – 177), **D_2_:** HISG; *Klebsiella pneumoniae HS11286*; motif range (156 – 177). **A_3_:** PLHDHIGA_04004; *Citrobacter werkmanii NIB003*; motif range (180-186), **B_3_:** ECS3215; *Escherichia coli O157H7 Sakai*; motif range (315-321), C3: STY2617; *Salmonella enterica typhi*; motif range (204-210), **D_3_:** PRMB; *Klebsiella pneumoniae HS11286*; motif range (204-210),A4: PLHDHIGA_02079; *Citrobacter werkmanii NIB003*; motif range (357-414), **B_4_:** TYRS; *Escherichia coli UMN026*; motif range (357-414), **C_4_:** Tyrs_2; *Salmonella enterica Typhi*; motif range (357-414), **D_4_:** TY1673; *Klebsiella pneumoniae ozaenae*; motif range (357-414),A5: PLHDHIGA_03404; *Citrobacter werkmanii NIB003*; motif range (138 - 555), B5: ASPS; *Escherichia coli O157H7* EDL933; motif range (138 – 555), **C_5_**: ASPS; *Salmonella enterica Typhi*; motif range (121 - 538), **D_5_:** STY2109; *Klebsiella aerogenes*; motif range (121 - 538).

**Table 5.** Function of motifs of major interacting proteins of *Citrobacter werkmanii NIB003*.

| Sequence ID (C. werkmanii NIB003) | Motif Name | Motif function | Motif Range | Similar Bacteria |
| --- | --- | --- | --- | --- |
| PLHDHIGA_00433 | ATP-PRTase | Facilitates the initial stage of histidine production in bacteria | 156 - 177 | *E. coli* O157H7 Sakai, *S. enterica* Typhi, *K. pneumoniae* HS11286 |
| PLHDHIGA_04004 | Dam methylase | DNA is modified by Dam methylase | 180 - 186 | *E. coli* O157H7 Sakai, *S. enterica* Typhi, *K. pneumoniae* HS11286 |
| PLHDHIGA_01430 | Aminoacyl-transfer RNA synthetases class-II family | Involves in the activation of amino acids and their subsequent transfer onto designated tRNA molecules. | 1 - 327 | *E. coli* O157H7 Sakai, *S. enterica* Typhi, *K. aerogenes* |
| PLHDHIGA_01431 | Tetratricopeptide repeat-like domain | peroxisomal protein transport and Cell cycle regulation | 15- 205 | *E. coli* O157H7 EDL933, *S. bongori*, *K. sp. RITPId* |
| PLHDHIGA_01946 | TGS domain; Aminoacyl-transfer RNA synthetases class-II family | Involved in rigorous response in bacteria | 1-61, 243- 534 | *E. coli* O157H7 Sakai, *S. enterica* arizonae, *K. pneumoniae* ozaena |
| PLHDHIGA_01118 | tRNA-binding domain | To maximize the catalytic rate of aminoacyl-tRNA synthetases | 575 - 677 | *E. coli* CFT073, *S. enterica* diarizonae, *K. quasipneumoniae* |
| PLHDHIGA_01872 | Aminoacyl-transfer RNA synthetases class-II family | Involves the activation of amino acids and their subsequent transfer onto designated tRNA molecules | 139 - 456 | *E. coli* O157H7 Sakai, *S. enterica* houtenae, *K. pneumoniae* HS11286 |
| PLHDHIGA_02079 | S4 RNA-binding domain; Aminoacyl-transfer RNA synthetases class-I | Identify intricate and distinctive three-dimensional characteristics inside extensively folded RNA molecules, such as rRNAs and tRNAs | 357- 414, 42 - 52 | *E. coli* UMN026, *S. enterica* Typhi, *K. pneumoniae* ozaenae |
| PLHDHIGA_03404 | Aminoacyl-transfer RNA synthetases class-II family | Involves the activation of amino acids and their subsequent transfer onto designated tRNA molecules. | 138 - 555 | *E. coli O157H7 EDL933*, *S. enterica Typhi*, *K. aerogenes* |
| PLHDHIGA_02178 | tRNA synthetases class I, catalytic domain | Synthesis of the aminoacyl-adenylate is under the responsibility of this enzyme | 3-313 | *E. coli O157H7 Sakai*, *S. enterica diarizonae*, *K. aerogenes* |
| PLHDHIGA_00328 | Aminoacyl-transfer RNA synthetases class-II | Plays a role in protein biosynthesis | 171 - 416 | *E. coli O157H7 Sakai*, *S. enterica diarizonae*, *K. oxytoca* |

### Intragenomic differences between *C. werkmanii* species

We investigated the intragenomic differences of *C. werkmanii* with two whole genomes (C*. werkmanii FDAARGOS_616, C. werkmanii LCU-V21*) from NCBI database, and compared them with our 11 major interacting proteins found from our own sequenced *C. werkmanii NIB003* species using Blastp. Despite having significant similarity, we found some substantial changes in our *C. werkmanii NIB003* genome. We showed the percentages of similarity of our *C. werkmanii NIB003* with those two bacterial strains (*C. werkmanii FDAARGOS_616*, *C. werkmanii LCU-V21*) (**Supplementary table 1**).

Additionally, we found three distinctive motifs in *C. werkmanii NIB003*, *C. werkmanii FDAARGOS_616, C. werkmanii LCU-V21* using Blastp and ScanProsite tools. The N6-adenine specific DNA methylase motif found in *C. werkmanii NIB003*, was absent in *C. werkmanii FDAARGOS_616*, *C. werkmanii LCU-V2*. In lieu of the N6-adenine specific DNA methylase motif, SAM-dependent methyltransferase RNAm(5)U-type domain was found in these two organism (*C. werkmanii FDAARGOS_616, C. werkmanii LCU-V21*) (**Supplementary table 1**).

### Molecular Dynamic Simulation

A root mean square deviation (RMSD) calculation is conducted to assess the stability of the system. The representation of PLHDHIGA (*C. werkmanii NIB003*, pink line), QET165917(C. werkmanii FDAARGOS_616, blue line), and MDO8232651 (*C. werkmanii LCU-V2*, green line) proteins has been illustrated in (**Fig. 7A)**. The proteins initially underwent a conformational change for 25 nanoseconds, after which they demonstrated stability. The RMSD value for the protein MDO8232651 was ∼0.5–0.6 nm, while the proteins PLHDHIGA and MDO8232651 exhibited an RMSD value of approximately 0.2-0.4 nm (**Fig. 7A**). As a result, the pattern of conformational change between MDO8232651 and the other two proteins (PLHDHIGA and MDO8232651) was significantly different. Moreover, the proteins PLHDHIGA and MDO8232651 showed similar patterns of conformational change.

**Figure 7.**
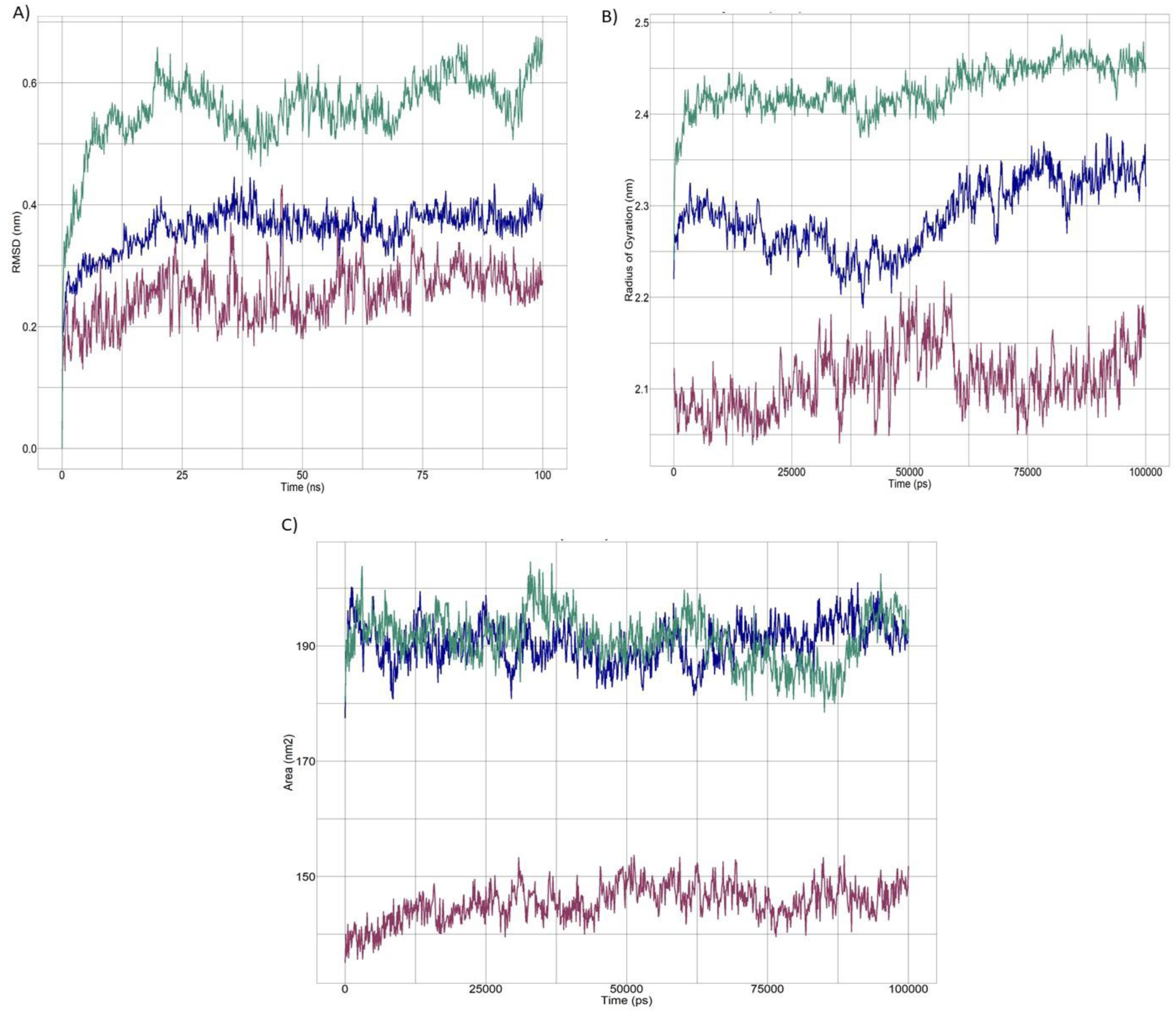
Root Mean Square Deviation (RMSD), Radius of gyration (Rg), Solvent Accessible Surface area (SASA) for 3 protein structures (PLHDHIGA_04004, QET165917, MDO8232651) of *Citrobacter werkmanii* species (*C. werkmanii NIB003*; *C. werkmanii FDAARGOS_616*; *C. werkmanii LCU-V2*). PLHDHIGA_04004 (*C. werkmanii NIB003)* in pink line, QET1659179 (*C. werkmanii FDAARGOS_616*) in blue line, MDO8232651(*C. werkmanii LCU-V2)* in green line.

The radius of gyration (Rg) calculates the degree of compactness. A steady radius of gyration indicates the protein is undergoing stable folding, whereas variations in the radius of gyration indicate the protein is unfolding. In contrast to the fluctuating patterns observed in the compactness of PLHDHIGA (pink line) and QET165917 (blue line), the compactness of MDO8232651 (green line) was relatively consistent. (**Fig. 7B**).

In molecular dynamics simulations, Solvent Accessible Surface Area (SASA) is utilized to predict the exposure of the hydrophobic protein core to solvents. Higher SASA values indicate that the majority of the protein is accessible to water, whereas lower values indicate that the majority of the protein is concealed inside the hydrophobic core. SASA values of the proteins PLHDHIGA (pink line), QET165917 (blue line), and MDO8232651 (green line) were stable throughout the simulation (**Fig. 7C**). The protein PLHDHIGA (pink line) showed a lower value of SASA compared to the other two proteins.

Root Mean Square Fluctuation (RMSF) is used to determine the regional flexibility of the protein. An amino acid’s positional flexibility increases with increasing RMSF. The mobility of PLHDHIGA (pink line), QET165917 (blue line), and MDO8232651 (green line) is displayed in (**Fig. 8D1-D3).** Two peaks were identified after the amino acid 200th for the protein PLHDHIGA, whereas peaks were observed from the amino acid 0th to the 100th for the proteins QET165917 and MDO8232651. Overall, the mobility of all the proteins has changed.

**Figure 8.**
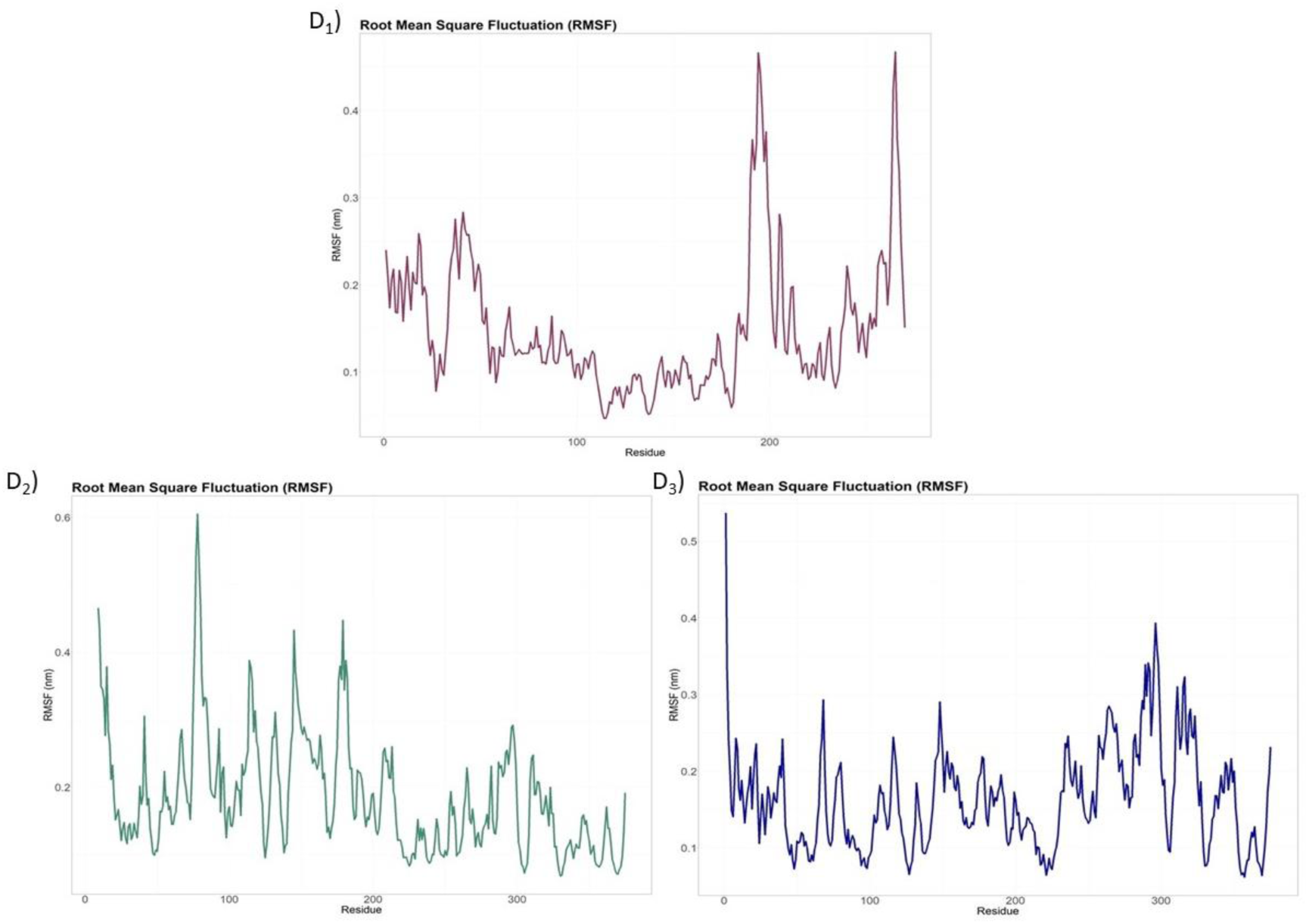
Root mean square fluctuation (RMSF) for 3 protein structures of *Citrobacter werkmanii* species (*C. werkmanii NIB003; C. werkmanii FDAARGOS_616; C. werkmanii LCU-V2*) of PLHDHIGA ((Pink line) and QET165917 (Blue line). MDO8232651 (Green line). *C. werkmanii FDAARGOS_616*; *C. werkmanii LCU-V2*). PLHDHIGA_04004 (*C. werkmanii NIB003)* in pink line, QET1659179 (*C. werkmanii FDAARGOS_616*) in blue line, MDO8232651(*C. werkmanii LCU-V2)* in green line.

## Discussion

*Citrobacter* species are opportunistic pathogens that lead to invasive illnesses such as diarrheal disorders. *C. werkmanii* has, however, only been reported in a small number of instances, mostly in Bangladesh. To gain a deeper understanding of this bacteria, a genome-wide study has been undertaken in order to investigate its genomic features. Firstly, *C. werkmanii* was identified using morphological observations, biochemical testing, and 16S rRNA sequencing on the isolated strain NIB003 (**Fig. 1**, **Fig. 2 and Table 1, Supplementary Fig. 1**). Few studies have identified the bacteria from the isolate, as biochemical tests and 16S rRNA gene sequences provide more reliable, accurate, and consistent bacterial identification (Y. Liu et al., 2022; Zhou et al., 2017). Following this, a complete genome of Bangladeshi isolates from diarrheal patient has been sequenced for the first time in Bangladesh. The genome ID is available at the National Centre for Biotechnology Information (NCBI) (**Fig. 3 and Table 2**). From the analysis of the complete genome of *C. werkmanii NIB003* genome, it was observed that a significant proportion of genes (170) were mainly connected with the synthesis of ABC transporters, which facilitate the movement of substances across the cell membranes of this bacteria by utilizing ATP hydrolysis as an energy source (**Fig. 4**). ABC transporters in bacteria are significant virulence factors due to their involvement in food acquisition and the secretion of toxins and antimicrobial compounds (Davidson & Chen, 2004). With the highest number of genes implicated in this particular pathway, it is highly probable that this bacterium exhibits resistance to many existing drugs (Akhtar & Turner, 2022). In addition, we also found a significant number of genes (45) involved in the flagellar assembly pathway (**Fig. 4**). The process of flagellar assembly is intricate and requires a significant amount of energy. Furthermore, flagellins play a significant role in the field of vaccine development as adjuvants, primarily because of their capacity to elicit robust immune responses (Nedeljković et al., 2021). In our isolated *C. werkmanii NIB003* strain, we identified a total of 43 genes involved in biofilm formation pathway, which is an indication of the virulence and pathogenicity of this bacteria (Stoodley et al., 2002). After rigorous investigations, we found two antibiotic resistance genes (blaCMY-98 and QnrB69_1), which confer resistance to beta-lactam drugs and fluoroquinolones respectively. Afterwards, we identified the 11 common interacting proteins in *C. werkmanii NIB003* proteome. Subsequently, we have constructed a protein-protein interaction network incorporating proteins from *C. werkmanii NIB003*, *E. coli*, *Klebsiella*, and *Salmonella* (**Fig. 5** and **Table 4**). In light of the functional role played by the domains and motifs of major interacting proteins in *C. werkmanii NIB003*, we conducted a comparative analysis of the domain present in *C. werkmanii NIB003* and those found in diarrheal pathogens including *Escherichia, Salmonella,* and *Klebsiella*. While there are notable similarities in protein structures and functions throughout all species, *C. werkmanii NIB003* exhibits different traits that may provide a promising avenue for innovative study centered on this bacterium (**Fig. 6**). The results of the phylogenetic study demonstrated a significant evolutionary relationship among *C. werkmanii NIB003*, *Salmonella*, *Escherichia*, and *Klebsiella*, as determined by domain and motif analyses (**Supplementary Fig. 2 and Supplementary Fig. 3**). In the genome comparison study, it was found that *C. werkmanii NIB003* had the most similar evolutionary relationships with *C. werkmanii FDAARGOS_616*, and *C. werkmanii LCU-V21*(**Supplementary Table 1**). However, it is worth noting that *C. werkmanii NIB003* possesses certain motifs, such as the t-RNA binding domain and N-6 Adenine specific DNA methylases, which are not present in both C*. werkmanii FDAARGOS_616* and *C. werkmanii LCU-V21*. Additional investigation is required in order to ascertain the underlying cause of this observed difference in functionality. There is a possibility that the occurrence of the phenomenon under consideration could be attributed to factors such as genetic mutations in the specific region.

The genetic changes were closely observed among the three genomes. Therefore, GROMACS (version 22.3) software was utilized to investigate the mutational impact of the proteins. The findings of our investigation reveal significant discrepancies in the structural modifications seen in the proteins PLHDHIGA (*C. werkmanii NIB003*), QET165917 (*C. werkmanii FDAARGOS_616*), and MDO8232651 (*C. werkmanii LCU-V21*).

The Root Mean Square Deviation (RMSD) (**Fig. 7A**) analysis revealed that the structure of PLHDHIGA remained roughly similar to that of QET165917, whereas MDO8232651 underwent a discernible structural alteration. Moreover, the analysis of the radius of gyration indicated that PLHDHIGA exhibited alterations in its compactness, similar to QET165917. In contrast, MDO8232651 constantly maintained a compact structure (**Fig. 7B**). The examination of Solvent Accessible Surface Area (SASA) revealed that the protein PLHDHIGA exhibited a comparatively lower SASA value in comparison to the other two proteins, suggesting variations in solvent exposure (**Fig. 7C**). The examination of Root Mean Square Fluctuation (RMSF) demonstrated the presence of diverse patterns of regional flexibility. Specifically, the protein PLHDHIGA exhibited two separate peaks, while QET165917 and MDO8232651 displayed peaks mostly in the early region (**Fig. 8 D1, D2, D3**). In general, the simulation outcomes offer valuable observations regarding the stability, conformational alterations, compactness, solvent exposure, and spatial flexibility of the protein-ligand complexes under investigation. These findings underscore the unique characteristics exhibited by PLHDHIGA when compared to MDO8232651 and QET165917.

Despite the whole genome sequencing of *C. werkmanii NIB003* and its genomic exploration could establish a robust platform and opportunity for therapeutic target and design, it still has some limitations due to methodological shortcomings such as sample size, extensive bioinformatics research and mice model study.

## Conclusion

The current study has been the first report in Bangladesh to sequence the complete genome of opportunistic *C. werkmanii* bacteria, which was further investigated for the genomic features. Subsequently, it becomes even more intriguing to explore when the findings indicate the distinct genetic makeup of its closest strain. Furthermore, the genetical association with the diarrheal pathogens in terms of their biological, molecular, and pathway might explore a new avenue in the study of the pathophysiology of the diarrheal diseases.

## Data Availability

The datasets that are displayed in this search can be accessed through online resources. Following are the names of the repository(s) and accession number(s) that can be found.

https://www.ncbi.nlm.nih.gov/nuccore/JAWLLA000000000

https://www.ncbi.nlm.nih.gov/bioproject/PRJNA1030290

https://www.ncbi.nlm.nih.gov/biosample/SAMN37903014

## Acknowledgements

The authors are grateful to microbial biotechnology division for providing the lab access during this project.

