## Supplementary file for "From Sequence to Significance: A Thorough Investigation of the Distinctive Genome Traits Uncovered in *C. werkmanii* strain NIB003"

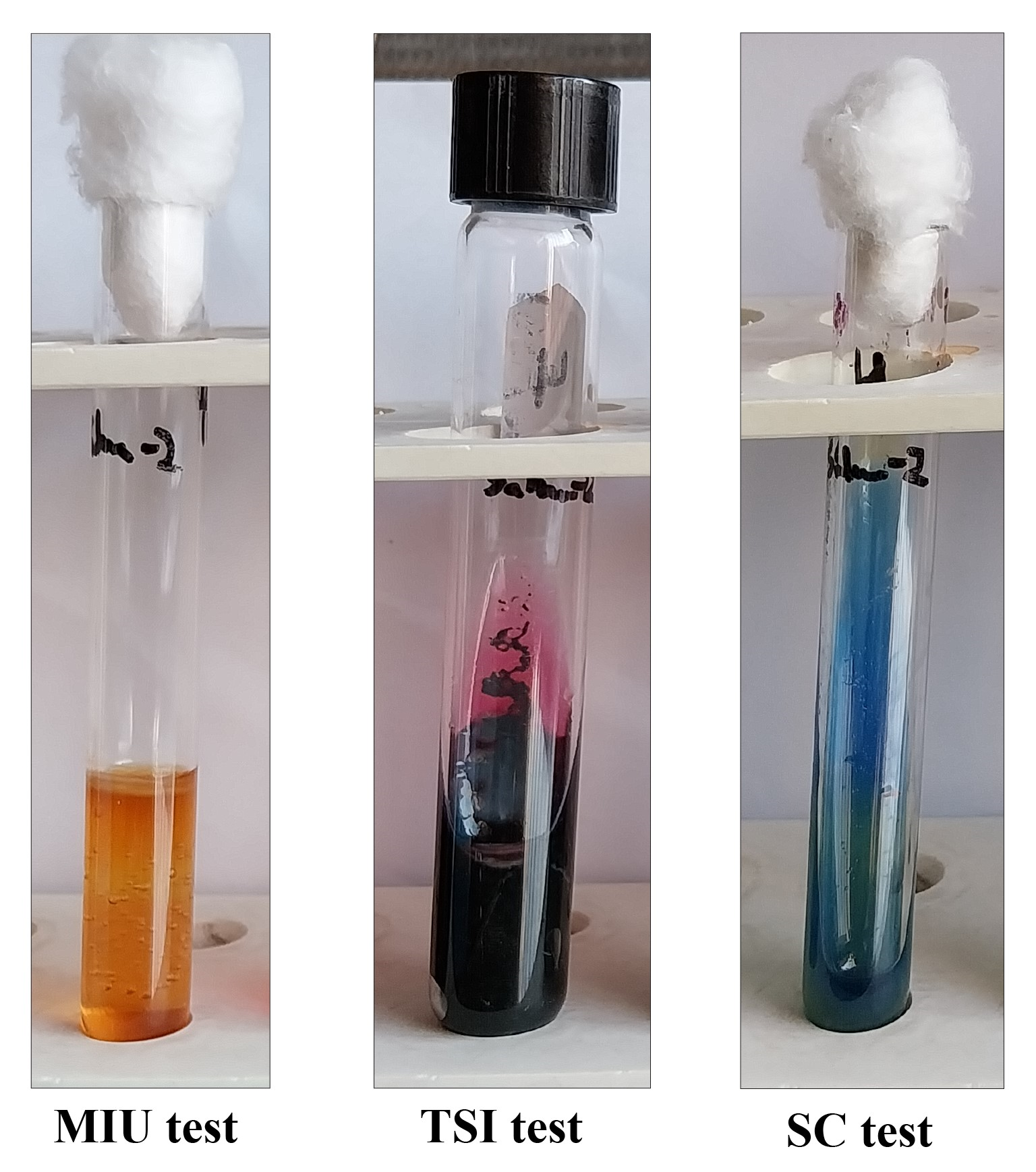

**Supplementary Figure 1** Different biochemical test (MIU test, TSI test, SC test) in test tube for Citrobacter werkmanii.

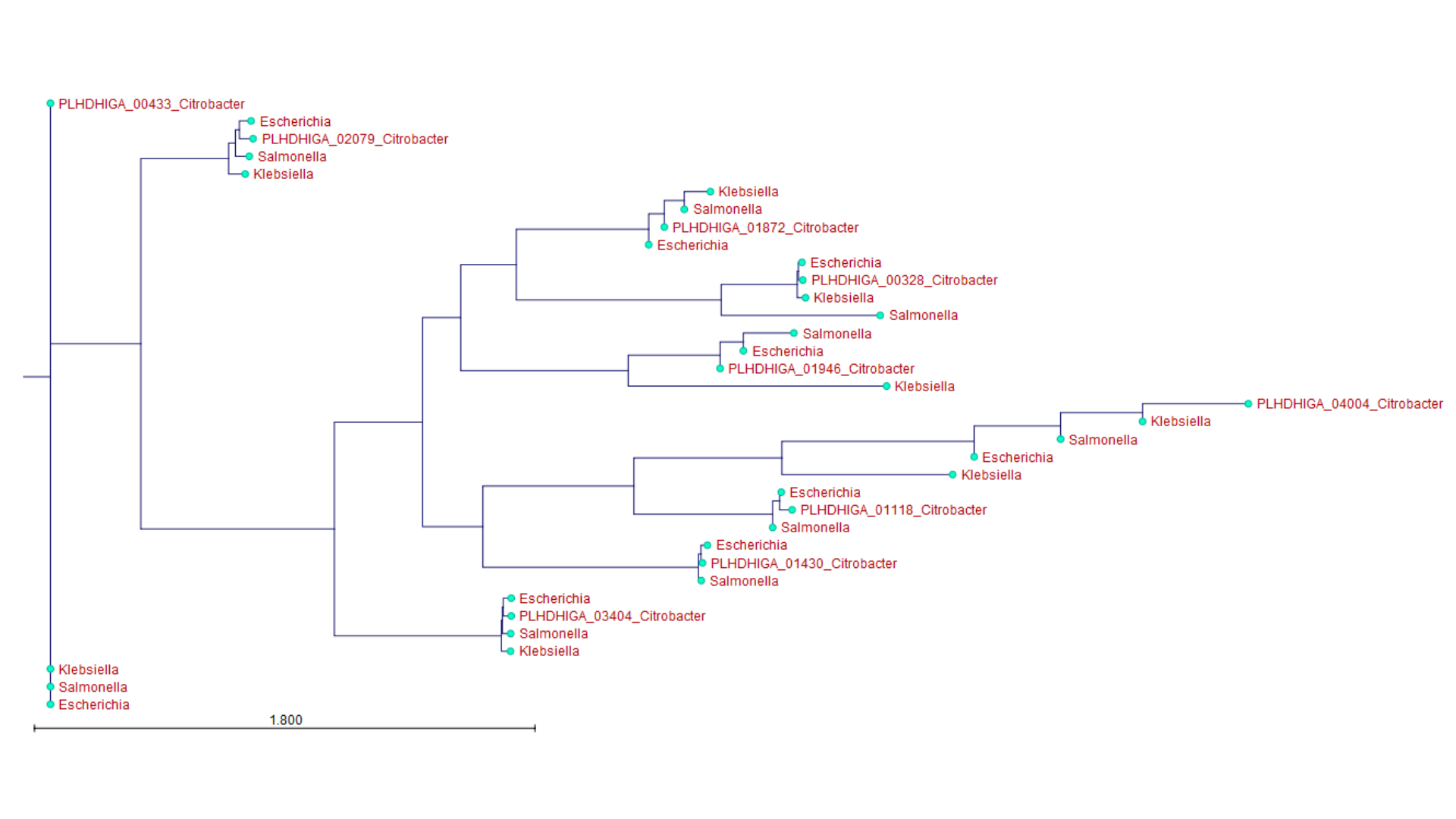
**Supplementary Figure 2** Phylogenetic trees of sequences from *Citrobacter werkmanii NIB003, Escherichia, Salmonella,* and *Klebsiella.*

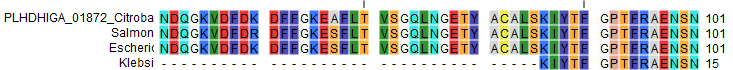

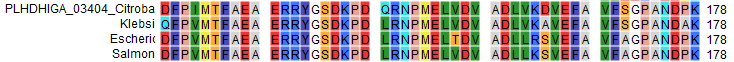

**Supplementary Figure 3** Multiple sequence alignment of sequences from *Citrobacter werkmanii NIB003, Escherichia, Salmonella,* and *Klebsiella.*

**Supplementary Table 1** Comparison of *Citrobacter werkmanii NIB003* with *C. werkmanii FDAARGOS_616* and *C. werkmanii LCU-V21*

| **Major interacting proteins (*Citrobacter werkmanii NIB003*)** | ***C. werkmanii FDAARGOS_616* (%similarity with major interacting proteins/genes)** | ***C. werkmanii LCU-V21* (%similarity with major interacting proteins/genes)** |
| --- | --- | --- |
| PLHDHIGA_01946  (Threonine-tRNA ligase) | QET65219.1  proS (62%) | MDO8235309.1  (54%) |
| PLHDHIGA_02079  (tyrosine--tRNA ligase) | QET66389.1  tyrS (100%) | MDO8233224.1  tyrS (100%) |
| PLHDHIGA_03404  (Aspartate-tRNA ligase) | QET67865.1  lysS (76%);  QET65963.1  asnS (58%);  QET64754.1  epmA (30%) | MDO8234936.1  lysS (76%);  MDO8232698.1  asnS (57%);  MDO8235803.1  epmA (30%); |
| PLHDHIGA_00328  (Serine-tRNA ligase) | QET65219.1  proS (37%);  QET66499.1  FOB24_05030 (15%) | MDO8235309.1  proS (37%);  MDO8233333.1  FOB24_05030 (15%) |
| PLHDHIGA_01872  (Asparagine-tRNA ligase) | QET67865.1  lysS (95%);  QET66861.1  aspS (77%);  QET64754.1  epmA (72%) | MDO8234936.1  lysS (95%);  MDO8233887.1  aspS (77%);  MDO8235803.1  epmA (72%) |
| PLHDHIGA_02178 (Cysteine-tRNA ligase) | QET65058.1  ileS (12%);  QET67123.1  metG (14%) | MDO8235125.1 tRNA-binding protein(15%)  MDO8232439.1  leuS (21%)  MDO8235477.1  ileS (7%)  MDO8232317.1  cysS (10%)  MDO8235675.1  valS (6%) |
| PLHDHIGA_01118 (methionine--tRNA ligase) | QET65675.1  leuS (21%);  QET65058.1  ileS (7%);  QET65561.1  cysS (10%);  QET64870.1  valS (6%) | MDO8235125.1 tRNA-binding protein (15%);  MDO8232439.1  leuS (21%);  MDO8235477.1  ileS (7%);  MDO8232317.1  cysS (10%);  MDO8235675.1 valS (6%) |
| PLHDHIGA_01431  (tetratricopeptide repeat protein) | QET67472.1 (tetratricopeptide repeat protein)100% | MDO8234584.1 YfgM family protein (100%) |
| PLHDHIGA_01430 (Histidine-tRNA ligase) | QET67473.1  hisS (100%) | MDO8234585.1  hisS (100%) |
| PLHDHIGA_04004  (Protein-(glutamine-N5) methyltransferase) | QET67325.1  uL3 MTase (69%);  QET65004.1  RsmC (29%);  QET66985.1  cbiT (22%);  QET65917.1  RlmC (25%) | MDO8234442.1  uL3 MTase (69%);  MDO8235534.1  RsmC(29%);  MDO8234072.1  cbiT (22%);  MDO8232651.1  RlmC (25%); |
| PLHDHIGA_00433(ATP phosphoribosyltransferase) | QET67025.1 ATP phosphoribosyltransferase (100%) | MDO8234114.1 ATP phosphoribosyltransferase(100%) |
